# Biochemical and structural insights into the activation of PBP1b by the essential domain of FtsN

**DOI:** 10.1101/2020.06.05.136150

**Authors:** Adrien Boes, Frederic Kerff, Raphael Herman, Thierry Touze, Eefjan Breukink, Mohammed Terrak

## Abstract

Peptidoglycan (PG) is an essential constituent of the bacterial cell wall. During cell division PG synthesis localizes at mid-cell under the control of a multiprotein complex, the divisome. In *Escherichia coli*, septal PG synthesis and cell constriction rely on the accumulation of FtsN at the division site. The region L75 to Q93 of FtsN (^E^FtsN) was shown to be essential and sufficient for its functioning *in vivo* but the specific target and the molecular mechanism remained unknown. Here, we show that ^E^FtsN binds specifically to the major PG synthase PBP1b and is sufficient to stimulate its GTase activity. We also report the crystal structure of PBP1b in complex with ^E^FtsN which provides structural insights into the mode of binding of ^E^FtsN at the junction between the GTase and UB2H domains of PBP1b. Interestingly, the mutations R141A/R397A of PBP1b, within the ^E^FtsN binding pocket, reduce the activation of PBP1b by FtsN. This mutant was unable to rescue Δ*ponB*-*ponA*^ts^ strain at nonpermissive temperature and induced a mild cell chaining phenotype and cell lysis. Altogether, the results show that PBP1b is a target of ^E^FtsN and suggest that binding of FtsN to PBP1b contributes to trigger septal PG synthesis and cell constriction.

## Introduction

Peptidoglycan (PG) is an essential constituent of the bacterial cell wall and a major antibacterial target; it surrounds the cytoplasmic membrane, determines the cell shape and protects the cell from rupture under internal osmotic pressure. The PG structure consists of glycan strands made of alternating β-1,4-linked *N*-acetylglucosamine (GlcNAc) and *N*-acetylmuramic acid (MurNAc) residues cross-linked by peptides ^1^. It is assembled using the lipid II precursor (undecaprenyl-pyrophosphoryl-MurNAc-(pentapeptide)-GlcNAc) by the GTase activities of the class A penicillin-binding proteins (aPBPs) and SEDS (shape, elongation, division, and sporulation) proteins; and cross-linked by the transpeptidase (TPase) activities of class A and class B PBPs (bPBPs) ^2–4^. The GTase and TPase activities are coupled within the same aPBPs and also with their bPBPs TPase partners (e.g., PBP1a-PBP2, PBP1b-PBP3) ^5^. Similarly, the GTase activity of the SEDS proteins are regulated by a cognate bPBP (RodA-PBP2 and FtsW-PBP3) ^3,6–8^. The activities of the PBPs was shown to be regulated within multiprotein complexes allowing a concerted PG synthesis and sacullus enlargement in line with the cell cycle progression ^9–12^.

Bacteria generally contain a set of PBPs with at least one bifunctional (GTase/TPase) PBP of class A and one monofunctional (TPase) PBP of class B ^2,13^. *E. coli* contains three aPBPs: PBP1a, 1b and 1c, and two bPBPs: PBP2 and PBP3, the latter being involved in cell elongation and division, respectively ^14^. PBP1a and PBP1b are major PG synthases and at least one of them is required for cell viability ^15^. Although the two PBPs are exchangeable, they likely play specific function during the cell cycle. PBP1a is mainly involved in cell elongation in partnership with PBP2, while PBP1b exhibits a preference for cell division in agreement with its enrichment at midcell during cell constriction ^12,16–18^. PBP1b has a modular structure, composed of an N-terminal tail and a transmembrane anchor followed by the GTase and TPase catalytic domains separated by the regulatory UB2H domain. PBP1b was shown *in vitro* to form a ternary complex with FtsW and PBP3 ^8^, which constitutes the septal synthase subcomplex of the divisome. This complex interacts with the divisome regulatory proteins FtsBLQ, FtsN and the outer-membrane lipoprotein LpoB ^9–11^. FtsN is the last essential protein that localizes at the division site; its accumulation, using a self-enhanced positive feedback mechanism, triggers cell constriction ^19^. LpoB binds to the UB2H domain of PBP1b and stimulates both of its catalytic activities ^20^. FtsN and LpoB bind simultaneously to PBP1b and synergistically stimulate the GTase activity of PBP1b ^12^. The GTase activity of PBP1b is repressed by FtsBLQ (via FtsL) during divisome assembly and the presence of FtsN and/or LpoB suppresses this inhibition ^9^. Moreover, FtsBLQ (via FtsQ) was also shown to inhibit the TPase activity of PBP3 but not that of PBP1b ^9^. On the other hand, CpoB interacts with TolA and both proteins bind to the PBP1b-LpoB complex, between the UB2H and TPase domains, thus inhibiting the TPase activity of PBP1b but not its GTase activity ^12^. This regulatory system blocks septal PG (sPG) synthesis thus, repressing cell division until the maturation of the divisome is signaled by the accumulation of FtsN, which stimulates PBP1b and counterbalances the inhibitory effect of FtsBLQ triggering sPG synthesis and the initiation of cell constriction.

From a structural point of view, FtsN is a bitopic membrane protein composed of a small cytoplasmic domain that interacts with FtsA at the 1C sub-domain, a transmembrane α-helix and a large periplasmic domain ^21,22^. The latter is further divided into three subdomains: a membrane-proximal portion containing three potential short α-helices, a glutamine rich central region and a PG binding SPOR (sporulation-related repeat) domain at the C-terminus which binds preferentially to glycan chains devoid of stem peptides ^23,24^. The region located around α-helix 2 (L75-Q94, ^E^FtsN) is essential for the function of FtsN ^19,25^.

In this work, we have used complementary techniques including fluorescence anisotropy binding assays, activity assays, cross-linking and x-ray crystallography, and obtained clear evidence that ^E^FtsN region is sufficient for direct binding to PBP1b and the stimulation of its GTase activity *in vitro.* Furthermore, we identified the binding site of ^E^FtsN within PBP1b being located between its GTase and UB2H domains and we showed the importance of this binding site in the function of PBP1b *in vivo*.

## Results

### ^E^FtsN is sufficient for the stimulation of the GTase activity of PBP1b

The stimulation of PBP1b *in vitro* by FtsN is well established ^5,26^. On the other hand, *in vivo*, a small periplasmic domain of FtsN (^E^FtsN: L75 to Q93) (**Fig 1A**) is required and sufficient for the function ^25^, but the molecular mechanism of PBP1b activation remains largely unknown. The first indication that ^E^FtsN was involved in PBP1b activation originates from the *in vitro* study of the FtsN mutant W83L, which has been shown to be crucial for FtsN activity *in vivo* ^25^. Indeed, FtsN^W83L^ was found to be less efficient in the stimulation of PBP1b activity and in the suppression of the inhibition on the GTase activity by FtsBLQ ^9^. In addition, the interaction between FtsN^W83L^ mutant and PBP1b was reduced compared to the wild-type protein. Based on these observations, we prepared the synthetic peptide (^E^FtsN: L75 to Q93) to test its effect on PBP1b. Interestingly, although a higher concentration than full length FtsN protein was required (50 μM vs ~1 μM), the peptide was able to stimulate the GTase activity of PBP1b (**Fig. 1B**), but has no effect on that of PBP1a (**Fig. 1C**), confirming that ^E^FtsN is directly responsible for PBP1b activation. This result is consistent with *in vivo* data showing that ^E^FtsN (GFP fusion) was functional only when overexpressed ^19,25^. When the activity of TPase domain of PBP1b was analyzed using S2d (a mimic of the D-Ala-D-Ala of the natural substrate) as substrate, the addition of FtsN had no effect (**Fig. 1D**), indicating that the protein only modulates the GTase activity but not the TPase activity of PBP1b.

**Figure 1.**
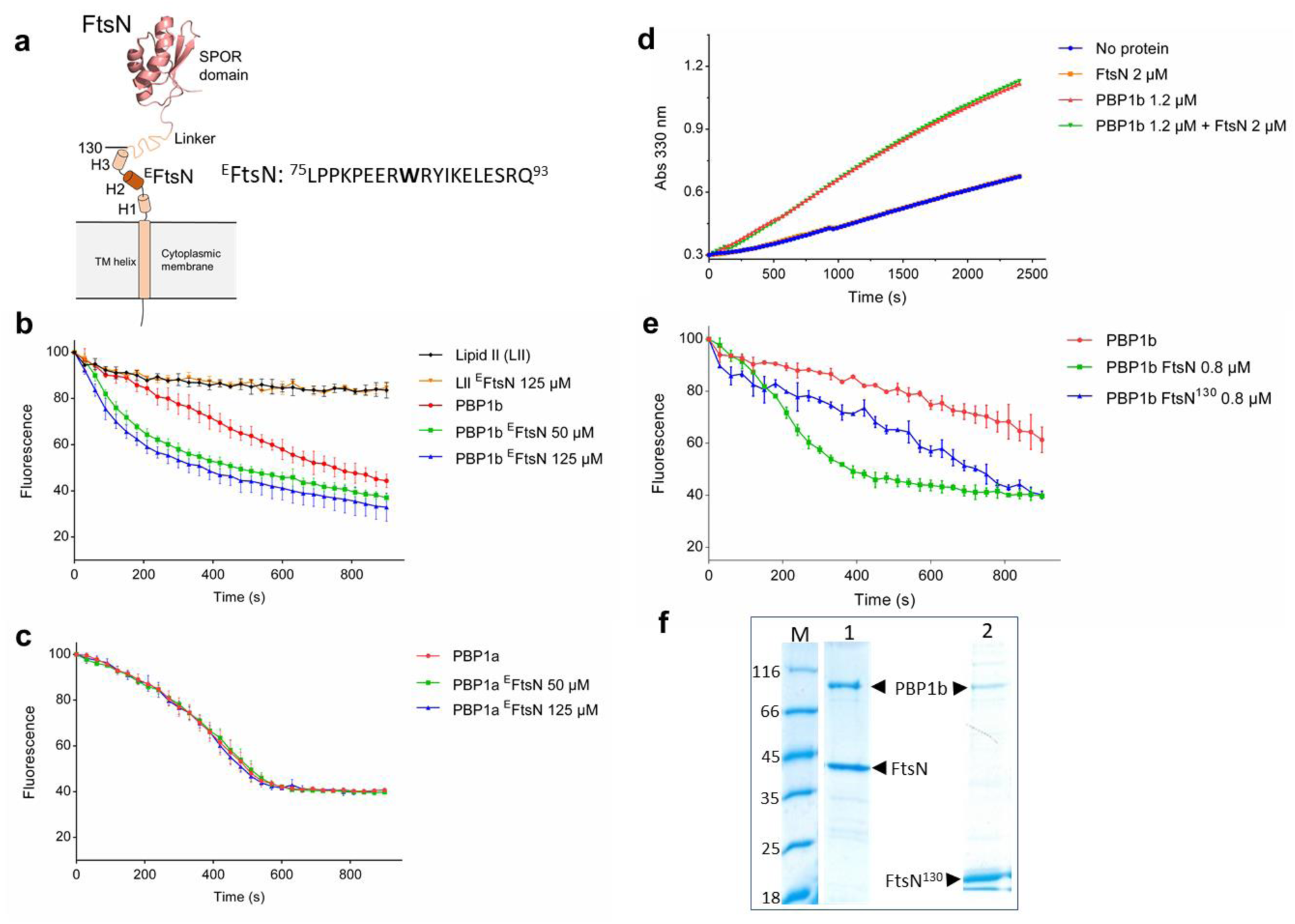
Activation of PBP1b GTase by ^E^FtsN and FtsN^130^. (A) Schematic representation of FtsN, the protein is composed of a short cytoplasmic tail and a transmembrane (TM) helix followed by 3 potential short helices (H1, H2 and H3), an unstructured Q-rich linker and a C-terminal SPOR domain (NMR structure 1UTA). The essential region of FtsN (^E^FtsN) correspond to the sequence L75-Q93 around H2. The construct FtsN^130^ (M1-R130) ends just after H3. The GTase activity of PBP1b and PBP1a was monitored by continuous fluorescence assay using dansyl-lipid II (LII) as substrate (B, C and E). The error bars represent the values as mean ± s.d. of three experiments. ^E^FtsN specifically stimulates the GTase activity of PBP1b (B) but has no effect on PBP1a (C). (D) The activity of the TPase domain of PBP1b was monitored by following the hydrolysis of S2d in the presence and absence of FtsN (a representative of 3 independent experiments). FtsN has no effect on the S2d hydrolysis by PBP1b (Abs 330 nm, absorbance at 330 nm). (E) Comparison of the stimulation of the GTase activity of PBP1b (1b) by FtsN and the truncated form FtsN^130^. (F) SDS-PAGE analysis of co-expression and co-purification of HisFtsN-PBP1b (line 1) and HisFtsN^130^-PBP1b (line 2). M, molecular mass marker.

We have also prepared a truncated variant of FtsN, called FtsN^130^ (encompassing residues M1 to R130) (**Fig 1. A, F**). This form, containing the three potential α-helices, was found to binds PBP1b and to stimulates its GTase activity to a lesser extent compared to full length protein, but was much more efficient in PBP1b activation than the ^E^FtsN peptide (**Fig. 1 E-F**). On the other hand, it was previously shown, using different variants of FtsN ^26^, including a soluble form lacking the cytoplasmic tail and the transmembrane (TM) anchor (sFtsN, Δ1-57), that multiple interaction sites contribute to the binding with PBP1b ^26^. The soluble form sFtsN displayed significantly reduced stimulatory effect on the PBP1b activity ^27^. Altogether, these data indicate that the N-terminal region, including the TM segment, is important for high affinity binding and efficient activation of PBP1b. In addition, the central Gln-rich region and SPOR domain seem to contribute to the optimal activation of PBP1b.

### ^E^FtsN interacts specifically with PBP1b but not with PBP3, FtsW-PBP3 or FtsBLQ

In order to measure the binding between ^E^FtsN and PBP1b or other divisome proteins a fluorescent peptide (FITC-K69-Q93), containing a FITC fluorophore attached at the N-terminus of the minimal sequence (L75-Q93) via six additional amino acids of FtsN (to avoid interference with binding), was prepared and used as a probe to develop a fluorescence anisotropy (FA) assay. Interaction studies with PBP1b show high increase of the FA signal and fitting of the graph allowed the determination of a dissociation constant *k*_*d*_ of 8.1 ± 1.7 μM (**Fig. 2A**). In contrast, no significant change in the FA signal was observed with PBP1a, PBP3, FtsW-PBP3 and FtsBLQ complexes (**Fig. 2A-B**), indicating that ^E^FtsN interacts specifically with PBP1b and excluding its direct interaction with FtsW-PBP3 or FtsBLQ as previously suggested ^25^. This result is consistent with the absence of strong interaction between FtsN and FtsBLQ and with the fact that FtsN has no effect on the activity of PBP3 ^9^.

**Figure 2.**
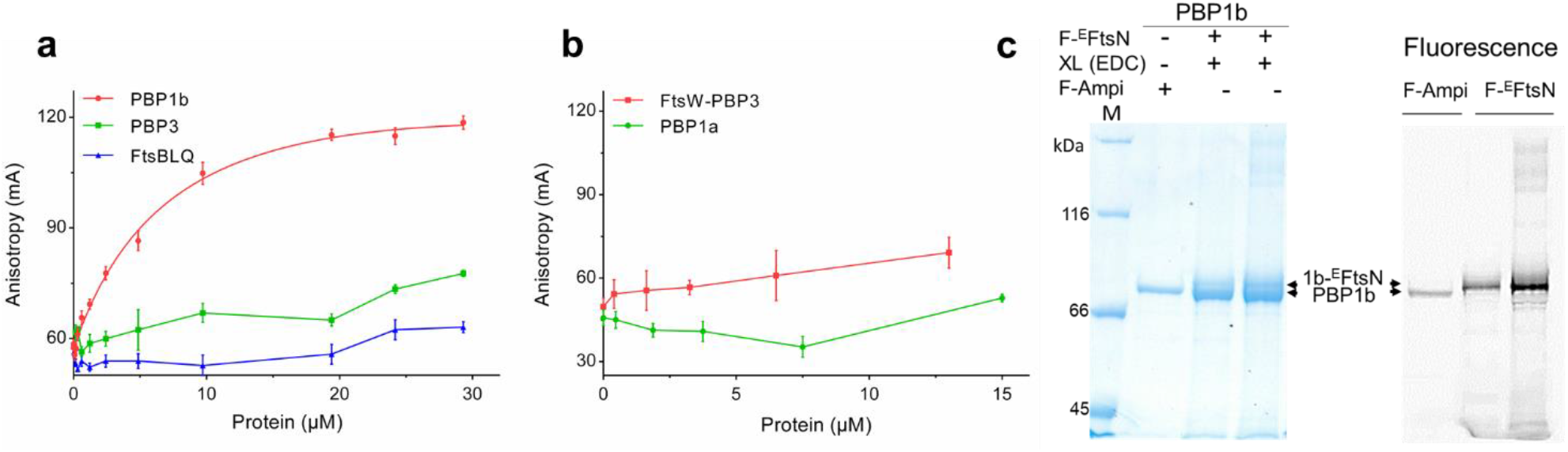
Binding assays between ^E^FtsN and PBP1b. **a**. Direct binding of fluorescent ^E^FtsN (F- ^E^FtsN) peptide to PBP1b using fluorescence anisotropy (FA) assay. FA (in mA units) is plotted as a function of protein concentrations. No significant binding is observed between the probe and PBP3, FtsBLQ complex, FtsW-PBP3 complex or PBP1a (**a** and **b**). The error bars represent the values as mean ± s.d. of three experiments. **c.** SDS-PAGE analysis of cross-linking (XL) adducts between PBP1b (1b) and F-^E^FtsN peptide (indicated by the upper arrows) using protein:cross-linker molar ratio of 1:1000 and 1:2000. Left, SDS-PAGE stained with Coomassie blue; right, fluorescence imaging. F-Ampi depicts PBP1b labelled with fluorescent ampicillin.

To further characterize the interaction between ^E^FtsN and PBP1b, we performed cross-linking experiments using the FITC-K69-Q93 peptide in order to facilitate the visualization of the adduct. The heterobifunctional cross-linking agent EDC (1-Ethyl-3-(3-dimethylaminopropyl)-carbodiimide), a zero-length cross-linker that couple carboxyl groups to primary amines, efficiently coupled PBP1b to the fluorescent peptide (**Fig. 2C**). SDS-PAGE analysis revealed either by Coomassie staining and by fluorescence labelling, shows a band slightly higher (PBP1b + peptide) than PBP1b alone labeled with fluorescent ampicillin. The band intensity increases when the concentration of the cross-linker was increased. This result further confirms that ^E^FtsN interacts with PBP1b.

### Crystal structure of PBP1b in complex with ^E^FtsN

Having strong evidence of the binding of ^E^FtsN to PBP1b, we performed co-crystallization assays between PBP1b (58-804), both in the presence and absence of moenomycin A, and purified ^E^FtsN (L75-Q93) peptide in order to identify the binding site of this domain on PBP1b. Crystals, which were only obtained in the presence of moenomycin A, belong to the space group P2_1_2_1_2 (**Table S1**), similar to those of previously reported for PBP1b structures ^28,29^. The structure was solved to 2.4 Å resolution by molecular replacement and showed an additional electron-density between the GTase and the UB2H domains (**Fig. 3A**). A 13 amino acid helix was fitted in this density and could be identified as residues P79-S91 from ^E^FtsN **(Fig 3A)**. This helix mostly interacts with the GTase domain and runs approximately parallel to its last α12 helix. Three hydrogen bounds are observed with PBP1b, between the side chain of W83 and the main chain carbonyl of Q384 (α11α12 loop, GTase domain), between the main chain carbonyl of L89 and side chain hydroxyl of T140 (α1β2 loop, UB2H domain), and between the E90 side chain and the R141 side chain (α1β2 loop, UB2H) as well as the L344 main chain (α9α10 loop, GTase) (**Fig. 3C-D**). In addition, Y85, I86 and L89 from ^E^FtsN form a hydrophobic cluster with L224 (α2), L344 (α9α10 loop), I386, L390 and L394 (α12) from the GTase domain (**Fig. 3C-D**).

**Figure 3.**
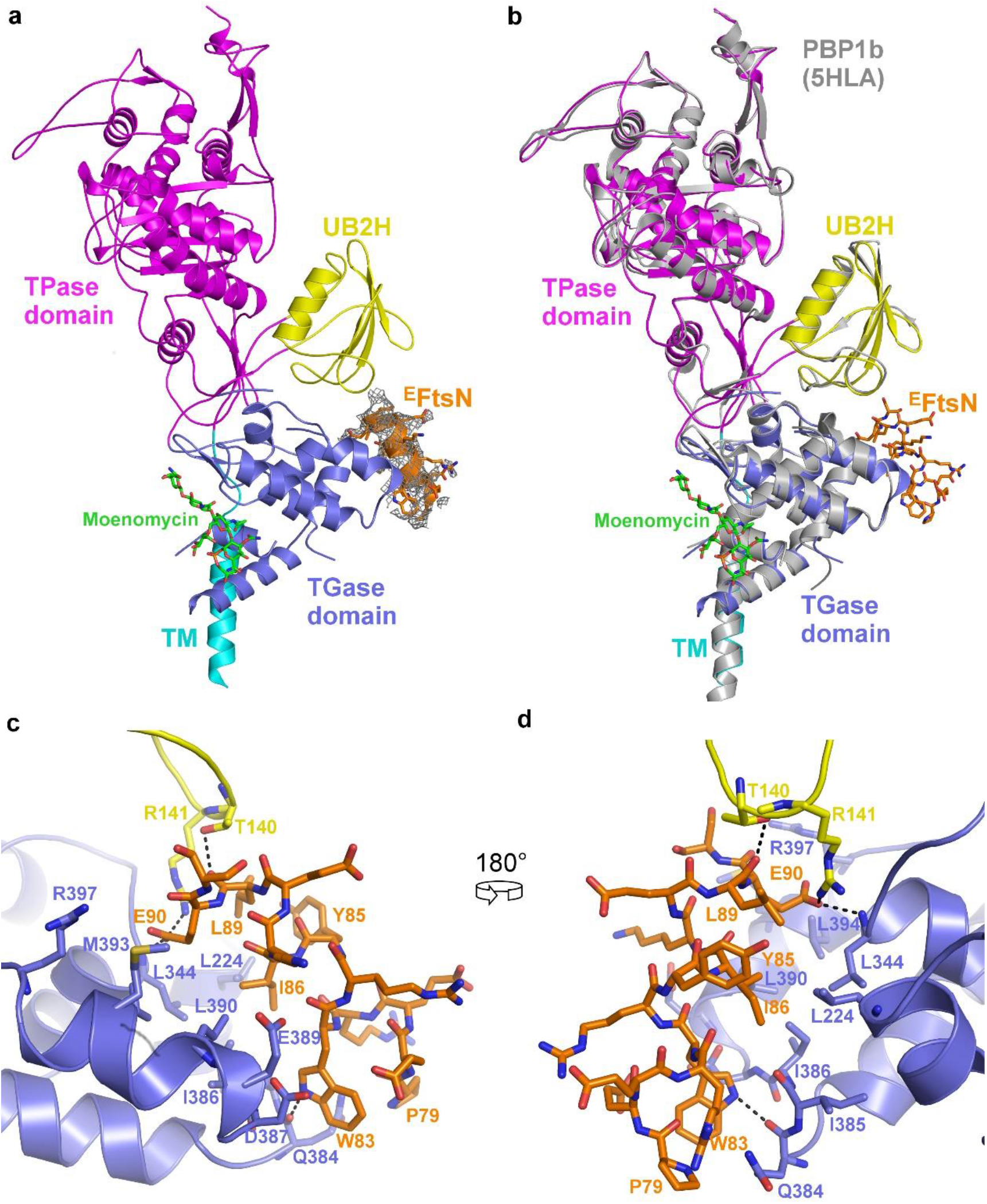
Crystal structure of PBP1b in complex with ^E^FtsN peptide. (A) Cartoon representation of the PBP1b-^E^FtsN structure with the 2*Fo* – *Fc* electron-density shown around ^E^FtsN (orange) at a 1 σ level in grey. The TM, UB2H, GTase, and TPase domains are shown with color code in cyan, yellow, blue, and magenta, respectively. This structure was obtained in the presence of moenomycin A shown in green sticks. (B) Superimposition of the obtained structure (same color code as in A) with that of that of published PBP1b structure (PDB code 5HLA) in grey. (C-D) details of the interactions between ^E^FtsN represented as orange sticks and the GTase (blue) and UB2H (yellow) domains of PBP1b presented in to orientations at 180° of each other. H-bonds are displayed as black dashed lines.

Compared with other PBP1b structures available, no major conformation change was observed upon ^E^FtsN binding with a rmsd of 0.85 Å over 685 common α carbons (PDB code 5HLA) (**Fig. 3B**). The differences are however larger in the GTase domain (rmsd 1.0 Å) than in the TPase (rmsd 0.56 Å) and the UB2H (rmsd 0.38 Å) domains. The GTase domain is also characterized by overall poorer electron densities compared to the rest of the protein as well as to other structures available. Of note, the crystals were only obtained in the presence of moenomycin A, therefore it is possible that this ligand may prevents the conformational change induced by the peptide. Nevertheless, we observed a significant destabilization of the GTase domain. This is materialized by a higher mean B factor for α carbons of the GTase domain (125 Å^2^) compared to other PBP1b structures (75 Å^2^ for 5HLA), while the TPase and UB2H domains are characterized by lower more similar values (50 Å^2^ and 48 Å^2^ respectively, versus 37 Å^2^ and 46 Å^2^ in the 5HLA structure). This could indicate that the conformational change responsible for the increased GTase activity may indeed be hindered by either the binding of moenomycin A required for crystallization, and/or the crystal packing, which strongly block the GTase domain in an inactive state while the peptide attempts to displace it to an active state.

### Mutations in the ^E^FtsN binding pocket reduce activation of PBP1b by FtsN and induce cell elongation and lysis

To confirm the importance of ^E^FtsN binding site in PBP1b activation, we addressed the roles of residues T140 and R141 from the UB2H domain side of the binding cavity that interact with ^E^FtsN via their side chains and the residue R397 from the GTase domain side of this cavity, which participate in the same positive patch as R141 and could contribute to the interaction with E90 of ^E^FtsN according to other PBP1b structures. All these residues were modified to Ala and single or double mutations were introduced in the *pon*B gene. The activity of the mutants was first evaluated by complementation experiments using *E. coli* EJ801 (Δ*ponB*-*ponA*^ts^) as a host strain (results in 10 g/L LB medium or low salt 0.5 g/L LB were comparable). The single mutants T140A, R141A, R397A and the double mutants T140A/R141A and T140A/R397A were able to restore the growth of the strain at the non-permissive temperature (42°C) (**Fig. 4A**) and the cells exhibited normal phenotype. In contrast, the double mutant PBP1b^R141A-R397A^ was unable to rescue the *E. coli* EJ801 at 42°C (**Fig. 4A**). The observation of the latter cells after 1 hour at the nonpermissive temperature showed mild elongated phenotype (L: 6.14 ± 1.84 μm *vs* 2.17 ± 0.46 μm) before lysis (**Fig. 4B-D**). Moreover, while the purified PBP1b^R141A-R397A^ mutant showed a minor difference in the GTase activity compared to the wild-type protein, its activation by FtsN was reduced by two-fold (**Fig. 4E**). These results indicate that, as a consequence of the mutations in ^E^FtsN binding site, the activation of PBP1b by FtsN was reduced causing defects in sPG integrity and cell division. This strongly confirms the link between PBP1b and FtsN during cell division and demonstrates the role of ^E^FtsN in the activation of its GTase activity.

**Figure 4.**
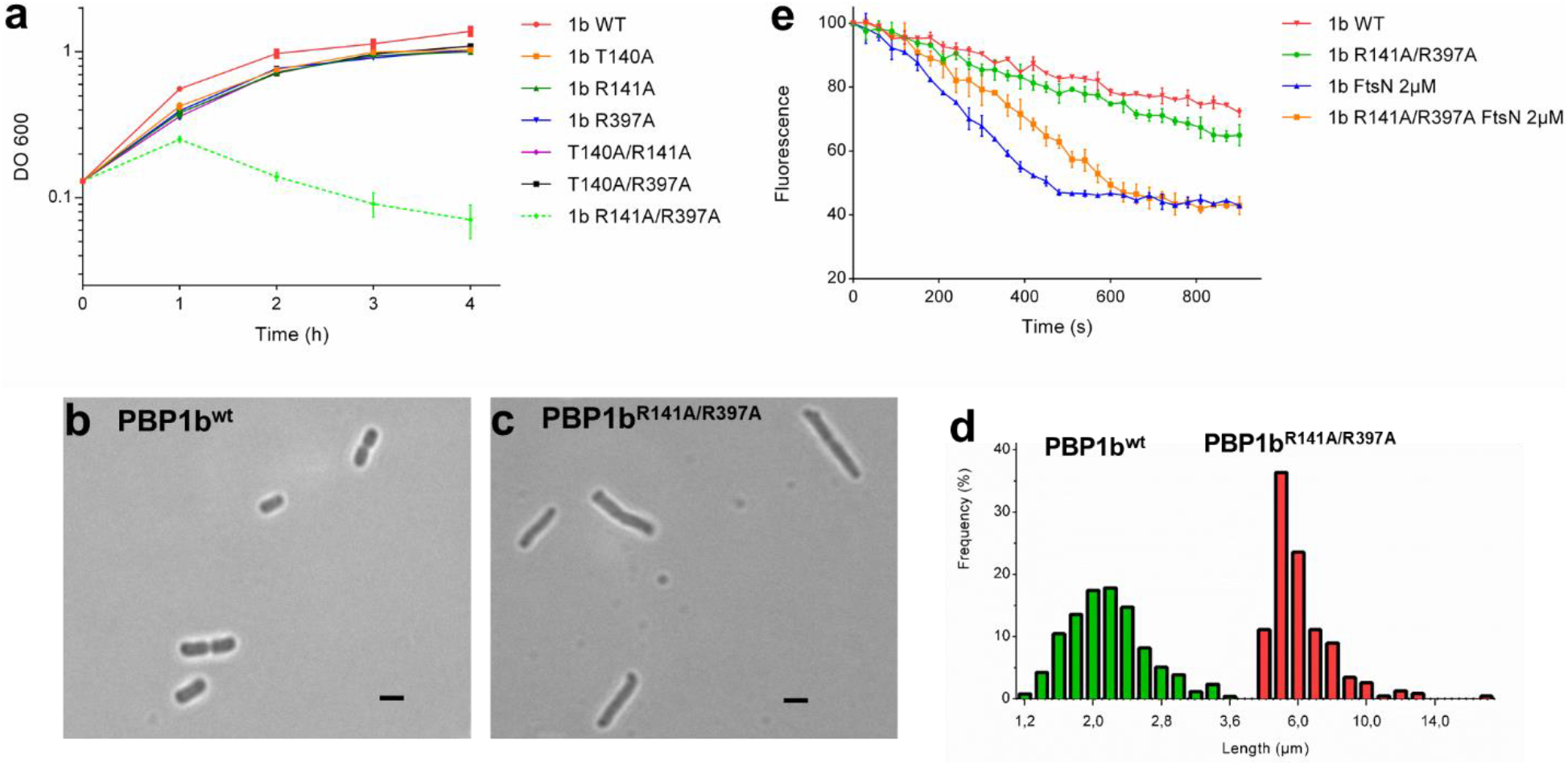
*In vitro* and *in vivo* activities of the PBP1b mutants. (A) complementation assays of *E. coli* EJ801 strain (*ΔponB-ponA*^*ts*^) with the plasmids carrying *ponB (*1b WT) gene or *ponB* mutants at non-permissive temperature (42°C). (B and C) microscopy analysis of the EJ801 cells expressing PBP1b^WT^ (B) or PBP1b^R141A/R397A^ (C) after 1h at 42°C. (D) Cell length statistics evaluation of EJ801 expressing PBP1b^WT^ (n = 250) or PBP1b^R141A/R397A^ (n = 250). (E) comparison of the GTase activation of PBP1b^WT^ (1b WT) and PBP1b^R141A/R397A^ by FtsN using continuous fluorescence assay. The error bars represent the values as mean ± s.d. of three experiments.

## Discussion

The mechanisms of *E. coli* septal PG synthesis and regulation are complex and involve several trans-envelope factors, with FtsA, FtsBLQ, FtsW, PBP3, PBP1b and FtsN playing direct and essential roles in these processes ^30–32^. Notably, PBP1b was shown to localize to the division site, to interact with several core divisome proteins and to be required for sPG synthesis ^8,9,12,18,26,33,34^. FtsN, on the one hand, and particularly its essential domain ^E^FtsN ^25^, plays a major role in the initiation of sPG synthesis that governs cell constriction, but what exactly ^E^FtsN does and its ultimate target remained unclear. On the other hand, biochemical data have shown that FtsN interacts with the major PG synthase PBP1b and stimulates its GTase activity ^5,26^. Using different techniques including FA binding assay, activity assay, cross-linking, X-ray crystallography, mutagenesis and complementation, we bring evidences that ^E^FtsN region is sufficient for direct binding to PBP1b and the stimulation of its GTase activity. Importantly, the crystal structure of the PBP1b-^E^FtsN complex allowed the identification of the binding site of ^E^FtsN being located between the GTase and UB2H domains of PBP1b (**Fig. 3**). The GTase domain accounts for most of the interaction with ^E^FtsN, but two residues (T140 and R141), in the α1β2 loop of the UB2H domain pointing towards the GTase domain, are also involved in the binding (**Fig. 3D**). This is consistent with the activation of the GTase but not the TPase activity of PBP1b by FtsN (**Fig. 1D-E)**. The binding site of ^E^FtsN is distinct from that of LpoB, which binds the exposed surface of UB2H domain ^20,35^. This is supported by the fact that FtsN and LpoB bind simultaneously to PBP1b and that the stimulation of the GTase activity of PBP1b by FtsN was synergistic with the stimulation by LpoB ^5^.

Furthermore, the structure reveals that the conserved residues W83, Y85 and L89 of ^E^FtsN, which were shown to be important for the *in vivo* function of the protein ^25^, together with I86 and E90 are the most critical for the interaction with PBP1b (**Fig. 3C-D**). Interestingly, the mutant FtsN^W83L^ was shown to exhibit reduced binding and activation of PBP1b ^9^. On the other hand, the double mutation R141A/R397A in the ^E^FtsN binding pocket reduces the activation of PBP1b by FtsN. In addition, this mutant was unable to rescue Δ*ponB*-*ponA*^ts^ strain at nonpermissive temperature and induced a mild cell chaining phenotype and cell lysis indicating sPG synthesis and division defects. These data suggest that the observed chaining phenotype is the result of the partial suppression of the ^E^FtsN binding site, which is a major regulatory hot spot for the stimulation of the GTase activity of PBP1b by FtsN during cell division. Altogether, the present results reveal that PBP1b, the major class A PBP, is a binding target of ^E^FtsN, and that PBP1b activation is mainly mediated by a direct interaction of ^E^FtsN with the GTase domain, although other parts of the protein are important for optimal function. However, our data do not support direct interaction between ^E^FtsN and FtsW-PBP3 or FtsBLQ, as previously proposed ^25^.

Overall, our results bridge the gap between the existing biochemical and *in vivo* data ^9,19,25,26,36^ and provide additional information to the general model of regulation of cell division in *E. coli* (**Fig. 5**). FtsN seems to play multiple and intricate functions (regulation of sPG synthesis and late hydrolysis) ^19,36^ that are coordinated with events on both sides of the cytoplasmic membrane, starting early in the cell cycle via its interaction with FtsA in the cytoplasm ^22,37–41^ and the gradual accumulation throughout the cell cycle ^42^, using a self-enhanced positive feedback mechanism ^19^. During divisome assembly, the GTase activity of PBP1b (as well as the TPase activity of PBP3) is kept inactive by the FtsBLQ complex until FtsN reaches a critical threshold which enables it to relieve this inhibition, probably by disrupting the interaction with FtsBLQ, and to restore PBP1b activity (via direct interaction with ^E^FtsN) ^9^ (**Fig. 5**). This, initiates sPG synthesis, probably in coordination with FtsW-PBP3 complex ^3,8,33^. sPG synthesis then recruits the amidases, that hydrolyze sPG and trigger cell constriction ^36^, this creates a high affinity binding substrate (denuded PG) of the SPOR domain allowing the recruitment of more FtsN and accelerates the constriction process ^23,24^.

**Figure 5.**
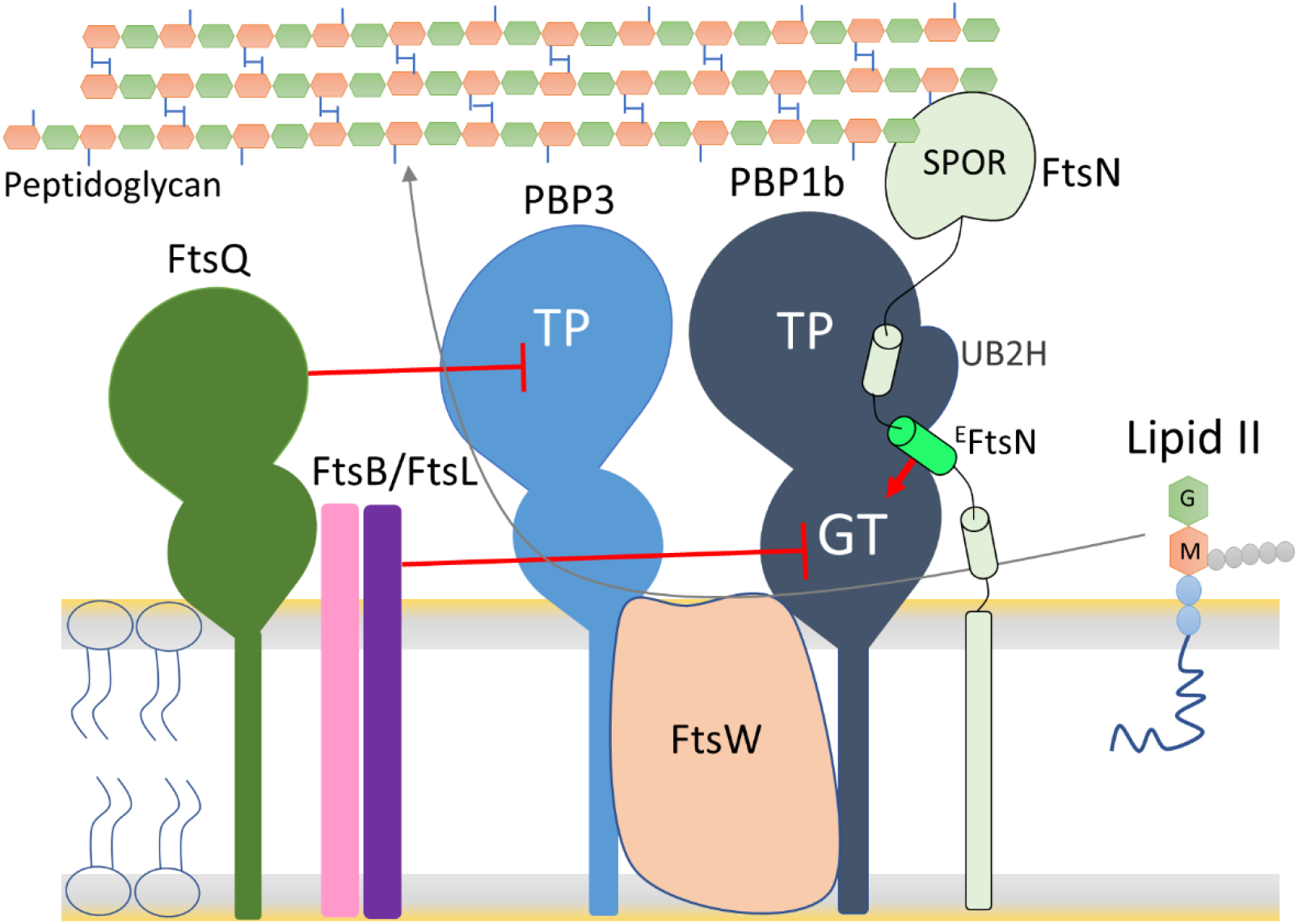
Model depicting the role of FtsN and its ^E^FtsN domain in the activation of sPG synthesis during cell division. During divisome assembly, the activity of the PBPs is inhibited by FtsBLQ complex, the accumulation of FtsN and its interaction with PBP1b brings the ^E^FtsN in contact with the GTase domain that activates sPG synthesis.

In *E. coli,* cell wall constriction depends mainly on sPG synthesis and not on FtsZ treadmilling ^34,43^. PBP1b was shown to be essential for sPG synthesis and cell constriction ^33,34^ and to exhibit two populations, a fast- and a slow-moving fractions ^44^. The slow-moving fraction represents the enzymes catalytically active in PG synthesis. FtsN also exhibits a dynamic behavior and its localization was shown to be spatially separated from constricting FtsZ-ring ^45^. As FtsN, a specific divisome protein, binds and activates PBP1b, it may play a role in the transition of PBP1b from the fast and low-activity state to the slow-moving processive enzyme at the division site.

PBP1b (and analogs in Gram-negative bacteria) is a major PG synthase required for mechanical stability of the PG sacculus and cell division which makes it an excellent target for new antibiotics ^33,44^. Perturbation of bacterial cell cycle (stimulation or repression) often leads to cell defects and eventually cell lysis. The identification of ^E^FtsN binding site could be useful in the search of allosteric modulators of PBP1b and related proteins for use as new antibiotics or in synergy with existing antibiotics against resistant strains.

## Materiel and Methods

### Bacterial strains, plasmids and growth conditions

#### Growth conditions

*E. coli* C43 (DE3) or Lemo21 (DE3) transformants were grown in Luria-Bertani (LB) or 2× YT medium supplemented with the appropriate antibiotic: ampicillin (100 μg/ml) (from MP Biomedicals), chloramphenicol (30 μg/ml) (from Sigma), or kanamycin (50 μg/ml) (from MP Biomedicals).

#### Reagents

Dansyl-lipid II was prepared as previously described ^46,47^. Fluorescein-labeled ampicillin was prepared as previously described ^48^. FITC labeled peptide (FITC-K69-Q93) KVTGNGLPPKPEERWRYIKELESRQ and unlabeled peptide (L75-Q93) LPPKPEERWRYIKELESRQ were purchased from Synpeptide co, (Shanghai China). They were solubilized in Tris-HCl 50 mM pH 8.0 and stored at −20°C until use. Moenomycin A was a gift from Aventis (Romainville, Paris).

### Plasmids construction

The complete list of the plasmids used in this study is given in **Table S2**. All point mutations were introduced using the Q5 site-directed mutagenesis kit (New England Biolabs). The primers used in this study are shown in **Table S3**; they were purchased from Eurogentec (Angleur, Belgium). Details for plasmid constructions are described in the supplementary information.

### Expression and purification of proteins

These proteins were purified as previously described: PBP1bγ ^49^, FtsN and FtsBLQ ^9^, PBP3 and FtsW-PBP3 ^8^, PBP1a ^50^. Further details for proteins purification are described in the supplementary information.

### Fluorescent anisotropy (FA) assay

FA experiments were performed to determine the binding affinity of the ^E^FtsN peptide, consisting of the fluorescein isothiocyanate (FITC)-K69-Q93 used as probe, to PBP1bγ. Serial dilutions of the proteins in their specific buffer were prepared in 384-well plates and the probe was added at 100 nM final concentration in a final volume of 30 μL. The mixtures were incubated for 30 min at 25°C and the FA signals were recorded using an Infinite® F Plex (Tecan, Männedorf, Switzerland) equipped with polarization filter with excitation wavelength at 485 nm and emission at 535 nm. FA values were calculated using the equations *FA* = (*I*_||_ – *G*·*I*_┴_)/*I*_||_ +2 *G*·*I*_┴_), where *I*_||_ is the fluorescence intensity of emitted light parallel to excitation, *I*_┴_ is the fluorescence intensity of emitted light perpendicular to excitation, and *G* is the correction factor that correct for instrument bias. The *G* factor is experimentally determined using the probe alone. For *K*_*d*_ determination, the fluorescence anisotropy data were analyzed by nonlinear curve fitting using GraphPad Prism 6.0 software as described ^51^.

### Continuous fluorescence GTase assay

The GTase activity assays with dansyl lipid II as substrate were performed in a medium binding black 96-well microplate (Greiner Bio One) as described ^52^. The samples contained 10 μM dansyl-lipid II, 50 mM HEPES-NaOH pH 7.5, 200 mM NaCl, 10 mM CaCl_2_, 0.085% of decyl-PEG, 20 % of dimethylsulfoxide (DMSO) and 1 unit of *N*-acetylmuramidase of *Streptomyces globisporus* (Sigma). The ^E^FtsN peptide L75-Q93 was solubilized in 50 mM Tris-HCl pH 8.0 and used at 50 μM and 125 μM concentrations. The proteins FtsN and FtsN^130^ were used at 0.8 μM. The reactions were initiated by the addition of 30 nM PBP1bγ (100 nM for PBP1a) and monitored by following the fluorescence decrease over 20 min at 30°C using an Infinite 200 PRO Microplate reader (Tecan, Männedorf, Switzerland) with excitation wavelength at 340 nm and emission at 520 nm.

#### Effect of FtsN on the hydrolysis of S2d by PBP1b

The activity of the TPase domain of PBP1b was measured in the presence of S2d (analog of the peptide moiety) as a mimic of donor substrates as previously described ^53,54^. The assay was performed in a UV-Star 96-well microplate (Greiner Bio One) at 37°C in the presence of 50 mM phosphate buffer pH 7.0, 2.0 mM S2d, 3.2 mM 4,4′-dithiodipyridine, and 1.2 μM PBP1b. The absorbance at 330 nm was monitored with an Infinite M200 Pro microplate reader (Tecan, Männedorf, Switzerland). FtsN was used at 2 μM to test its effect on the PBP1b activity. The experiments were repeated three times with reproducible results.

#### Cross-linking experiments

PBP1bγ (10 μM) was incubated with a 10-fold excess of FITC-^E^FtsN peptide and two conditions of the heterobifunctional crosslinker EDC (1-ethyl-3-(3-dimethylaminopropyl) carbodiimide hydrochloride) (protein:EDC molar ratio 1:1000 and 1:2000) in buffer 50 mM HEPES-NaOH pH 7,5, 0.3 mM NaCl, 0.7% CHAPS. The mixture was incubated for 2 h at room temperature and analyzed by SDS-PAGE followed by fluorescence imaging using a Typhoon Trio+ imager and Image Quant TL software (GE Healthcare).

### Crystallization, data collection, and structure determination

Crystallization was carried out using the hanging drop vapor diffusion method at 20°C. The PBP1b (K58-S804) concentration was 20 mg/ml in the buffer solution 20 mM Tris-HCl pH 8.0, 0.3 M NaCl and 4.5 mM DM and contained 1:1 molar ratio of moenomycin A and a synthetic peptide corresponding to ^E^FtsN (LPPKPEERWRYIKELESRQ) in a 3:1 molar ratio. The crystals were grown in drops made of 2 μl of protein solution and 2 μl of precipitant solution containing 0.1 M (NH4)_2_SO_4_, 0.3 M sodium Formate, 0.1 M Tris-HCl pH 7.8, 3% w/v low molecular-weight poly-γ-glutamic acids (PGA-LM) and 20 % v/v PEG 550 MME. The cryoprotectant solution was made of 22 % w/v PEG 6000 and 30 % v/v PEG 400. The diffraction data were collected on the Proxima 1 beamline of the Soleil synchrotron (Paris-Saclay). The data were indexed, integrated and scaled using XDS and reached a 2.4 Å resolution ^55^. Because of the anisotropy of the data, additional processing was carried out with Staraniso ^56^. The three axes of the ellipsoid used to cutoff the resolution were 2.3 Å, 4.3 Å and 2.3 Å. The crystal used belongs to the orthorhombic P21212 space group. The structure was solved by molecular replacement with Phaser ^57^ using the PBP1b structure of PBD code 3VMA as search model ^28^. The refinement and model building cycles were respectively performed with buster (BUSTER version 2.10.2. Cambridge, United Kingdom: Global Phasing Ltd.) and Coot ^58^. A summary of the relevant statistics of the data collection and refinement is given in **Table S1**. The figures were prepared using PyMOL (The PyMOL Molecular Graphics System, Version 1.7.4.3 Enhanced for Mac OS X, Schrödinger, LLC.). The coordinates and structure factors of the structure have been deposited in the Protein Data Bank with the PDB ID codes 6YN0.

#### Complementation *assay*

To test the activity of the PBP1b mutants *in vivo*, we tested their ability to complement *E. coli* strain EJ801 which lack PBP1b and has a thermosensitive PBP1a ^59^. EJ801 cells were transformed with the corresponding plasmids and grown in LB at 30 °C with 50 μg/ml kanamycine. The preculture was then diluted in 30 ml of fresh LB (10 g/ml NaCl) or LB with 0.5 g/ml NaCl medium to an OD_600nm_ value of 0.04 and grown at 30°C until DO_600_ 0.1–0.3, and the absorbance of the culture was monitored for 4 h at 42°C.

#### Microscopy and image analysis

Samples were taken from the complementation cultures (42°C) at different time (1h intervals) and the cells were fixed as described ^60^. Photographs were taken with a cooled AxioCam MRm (Zeiss) mounted on a Zeiss Axio Imager.Z1 microscope, and images were acquired in phase-contrast mode using the AxioVision Rel. 4.5 (Zeiss) software. The cell morphologies analysis were determined using ImageJ software (https://imagej.nih.gov/ij/) running under plugin ObjectJ (https://sils.fnwi.uva.nl/bcb/objectj/).

## Acknowledgements

This work was supported by the “Fonds de la Recherche Scientifique” CDR J.0030.18. MT and FK are research associates of the FRS_FNRS (Brussels, Belgium), AB is supported by FRIA 1.E.038.17 (Fonds pour la formation à la Recherche dans l’Industrie et dans l’Agriculture) fellowship of the FRS_FNRS. The assistance and support of beamline proxima 1 scientists at the Soleil synchrotron are acknowledged.

## Author contributions

AB, TT, RH, FK, EB performed research analyzed the data. MT designed the experiments, analyzed the data and wrote the manuscript with input from all the authors.

