## Supplementary material for "Biochemical and structural insights into the activation of PBP1b by the essential domain of FtsN"

**Running title:** Interactions and activities of PBP1b/ FtsN

**Keywords:** Peptidoglycan; divisome, PBP1b, FtsN, Lipid II.

### Plasmids construction

**pET28a-PBP1b(K58-S804).** The *E. coli pon*B gene encoding the truncated form of PBP1b corresponding to amino acids K58–S804 was amplified by PCR from pDML924 plasmid using the primers 1B_K58 and 1B_S804 (**Table S3**). PCR product was digested by NdeI/XhoI and ligated into the plasmid pET-28a between the corresponding sites. The resulting plasmid pET28a-PBP1b(K58-S804) encodes for PBP1b (K58-S804) polypeptide with an N-terminal His-tag followed by a thrombin cleavage site.

**pET22b-*fts*N$^{130}$His.** The *fts*N1-130 fragment was amplified by PCR from pDML2032 (**Table S2**) using the primers N_M1/pET22b and N_R130/pET22b (**Table S3**), digested by NdeI/XhoI and cloned into the plasmid pET22b between the corresponding sites. The encoded protein contains a His-tag at the C-terminus.

**pDuet-His*fts*N$^{130}$-*pon*B**. *fts*N1-130 fragment was amplified by PCR from pDML2032 using N_M1/pDuet and N_R130/pDuet (**Table S3**) and inserted in the MCS1 of the pETDuet1-HisFtsN-PBP1b plasmid between BamHI/ HindIII to replace the full-length *fts*N gene (**Table S2**). FtsN$^{130}$ contains a His-tag at the N-terminus.

### Expression and purification of proteins

**PBP1b(58-S804).** C43(DE3) cells transformed with pET28-PBP1b(K58-S804) were grown in Luria-Bertani (LB) with kanamycin (50 µg/ml) at 37°C to an $A_{600nm}$ of 0.8. Then, the expression was induced for 3.5 hours by addition of 0.5 mM isopropyl β-D-1-thiogalactopyranoside (IPTG). Cells were collected by centrifugation at 4000 × *g* for 20 minutes at 15°C and resuspended in a buffer containing 20 mM Tris-HCl pH 8.0, 300 mM NaCl and EDTA-free protease inhibitor Cocktail (Roche). The bacterial cells were lysed by three passages through a cell homogeneizer (Emulsiflex C3 Avestin). After centrifugation at 4000 × *g* for 20 minutes at 4°C, the supernatant was recovered and was spun down at 150,000 × g for 1 hour at 4°C and then the membranes were solubilized in 20 mM Tris-HCl pH 8.0, 500 mM NaCl, 10% v/v glycerol, 40 mM *n*-dodecyl-β-D-maltopyranoside (DDM; Inalco) and complete EDTA-free protease inhibitors. The mixture was incubated for 1 hour at room temperature under gentle agitation followed by centrifugation at 150,000 × g for 1 hour at 4°C. The supernatant containing the solubilized membrane proteins was loaded onto a HisTrap column (GE HealthCare) conditioned in buffer A (20 mM Tris-HCl pH 8.0, 300 mM NaCl, 1 mM DDM). After a wash with buffer A supplemented with 100 mM imidazole, the protein was eluted using a linear gradient of imidazole from 100–500 mM. The pure fractions were pooled and desalted on a G25 Sephadex column (GE HealthCare). The N-terminal His-tag was cleaved by the addition of 5 unit of bovine α-thrombin (Sigma**)** per mg of PBP1b during overnight incubation at room temperature. The sample was loaded again onto a HisTrap column conditioned in buffer A, the protein without His- tag was recovered in the flow through and during washing step with buffer A. The protein was concentrated using a 50-kDa cutoff Amicon centrifugation unit. PBP1b was further purified on a Superdex 200 Increase 10/300 GL (GE Healthcare) conditioned with 20 mM Tris-HCl pH 8.0, 300 mM NaCl, 4.5 mM *n*-decyl-β-D-maltopyranoside (DM; Anatrace). The protein was finally concentrated until 20 mg/ml and used for crystallization.

**FtsN$^{130}$ and PBP1b-FtsN$^{130}$ complex.** C43(DE3) cells transformed with pET22b-*fts*N$^{130}$His or pDuet-His*fts*N$^{130}$-*pon*B were grown in Luria-Bertani (LB) with ampicillin (100 µg/ml) at 37°C to an $A_{600nm}$ of 0.8. Then expression was induced for 3.5 hours by addition of 0.5 mM IPTG. Cells were collected by centrifugation at 4000 × *g* for 20 minutes at 15°C and resuspended in a buffer containing 20 mM Tris-HCl pH 8.0, 300 mM NaCl and EDTA-free protease inhibitor Cocktail (Roche). The cells were lysed by three passages through a cell homogeneizer (Emulsiflex C3 Avestin®). After centrifugation at 4000 × *g* for 20 minutes at 4°C, the supernatant was recovered and was spun down at 150,000 × g for 1 hour at 4°C and then the membranes were solubilized in 25 mM Tris-HCl pH 8.0, 500 mM NaCl, 10% v/v glycerol, 40 mM DDM (Inalco) and complete EDTA-free protease inhibitors. The mixture was incubated for 1 hour at room temperature under gentle agitation followed by centrifugation at 150,000 × g for 1 hour at 4°C. The supernatant containing the solubilized membrane proteins was loaded onto a HisTrap column (GE HealthCare) conditioned in buffer B (25 mM Tris-HCl pH 7.5, 500 mM NaCl, 4 mM DDM). After a wash with buffer B supplemented with 100 mM imidazole, the proteins were eluted using a linear gradient of imidazole from 100–500 mM. The pure fractions were pooled and desalted on a G25 Sephadex column (GE HealthCare).

**Table S1. X-ray crystallographic data collection and refinement statistics of E coli PBP1b in complex with $^{E}$FtsN**

| | PBP1b-$^{E}$FtsN |
|---|---|
| **Data Collection:** | |
| Space group | $P2_12_12$ |
| a, b, c (Å) | 63.1, 283.0, 62.7 |
| α, β, γ (°) | 90, 90, 90 |
| Resolution range (Å)[a] | 47.2 – 2.4 (2.51 – 2.4) |
| <I>/<σI> [a] | 8.1 (1.5) |
| Completeness elliptical (%)[a] | 95.5 (85.9) |
| Completeness spherical (%)[a] | 57.1 (22.3) |
| Redundancy [a] | 8.5 (7.2) |
| **Refinement:** | |
| Resolution range (Å) | 47.2 – 2.4 |
| No. of unique reflections | 25830 |
| R work (%) | 22.5 |
| R free (%) | 25.6 |
| No. atoms | |
| Protein | 5525 |
| Ligands | 77 |
| Water | 55 |
| RMS deviations from ideal stereochemistry | |
| Bond lengths (Å) | 0.01 |
| Bond angles (°) | 1.5 |
| Mean B factor (Å$^2$) | |
| Protein | 78.2 |
| Ligands | 150 |
| Water | 36.1 |
| Ramachandran plot: | |
| Favoured region (%) | 94.2 |
| Allowed regions (%) | 5.4 |
| Outlier regions (%) | 0.4 |

**Table S2. Plasmids used in this study**

| Plasmid | Description | Reference |
|---|---|---|
| pDML924 | His-PBP1bγ | [1] |
| pDML924 (T140A) | His-PBP1bγ (T140A) | This work |
| pDML924 (R141A) | His-PBP1bγ (R141A) | This work |
| pDML924 (R397A) | His-PBP1bγ (R397A) | This work |
| pDML924 (T140A/R141A) | His-PBP1bγ (T140A/R141A) | This work |
| pDML924 (T140A/R397A) | His-PBP1bγ (T140A/R397A) | This work |
| pDML924 (R141A/R397A) | His-PBP1bγ (R141A/R397A) | This work |
| pET28-*pon*B(K58-S804) | His-PBP1b(K58-S804) | This work |
| pDML2032 | FtsN-His | [2] |
| pET22b-*fts*N130His | FtsN130-His | This work |
| pTK1A | His-PBP1A | [3] |
| pRSF-His*fts*BL$^*$Q | His-FtsB/FtsL$^*$/FtsQ | [4] |
| pDML2494 | His-PBP3 | [5] |
| pDuet-His*fts*N-*pon*B | His-FtsN / PBP1b | [4] |
| pDuet-His*fts*N130-*pon*B | His-FtsN130 / PBP1b | This work |

**Table S3: Oligonucleotides used in this study**

| Oligonucleotide name | Sequence 5'→ 3' |
|---|---|
| 1B_K58 | GATACGCATCTCGAGTTATGACGGCTGCTGCTGCATCTC |
| 1B_S804 | GATACGCATCTCGAGTTATGACGGCTGCTGCTGCATCTC |
| N_M1/pET22 | TATACATATGGCACAACGAGATTATG |
| N_R130/pET22 | TATACTCGAGGCGCATATCAGCCTGC |
| N_M1/pDuet | TATAGGATCCTGCACAACGAGATTATGTACG |
| N_R130/pDuet | TATAAAGCTTTTAGCGCATATCAGCCTGC |
| 1B_T140A_F | GTCGAAAATGGCGCGTCCTGGCG |
| 1B_T140A_R | ACCTGACGATACTGGGTC |
| 1B_R141A_F | GAAAATGACCGCGCCTGGCGAATTTACC |
| 1B_R141A_R | GACACCTGACGATACTGG |
| 1B_ R397A_F | GTTGAGTGCCGCGCCGCTGGGGG |
| 1B_ R397A_R | ATGTCATAGAGTTCTTGATCAATAATC |
| 1B_T140A/R141A_F | GTCGAAAATGGCGGCGCCTGGCGAATTTACC |
| 1B_T140A/R141A_R | ACCTGACGATACTGGGTC |
